# Data Note: Monash DaCRA fPET-fMRI: A DAtaset for Comparison of Radiotracer Administration for high temporal resolution functional FDG-PET

**DOI:** 10.1101/2021.08.02.454708

**Authors:** Sharna D Jamadar, Emma X Liang, Shenjun Zhong, Phillip GD Ward, Alexandra Carey, Richard McIntyre, Zhaolin Chen, Gary F Egan

**Author notes:** **Corresponding Author:** Sharna D Jamadar, PhD, 770 Blackburn Rd, Melbourne VIC 3800, Australia.

## Abstract

**Background:** ‘Functional’ [18F]-fluorodeoxyglucose positron emission tomography (FDG-*f*PET) is a new approach for measuring glucose uptake in the human brain. The goal of FDG-fPET is to maintain a constant plasma supply of radioactive FDG in order to track, with high temporal resolution, the dynamic uptake of glucose during neuronal activity that occurs in response to a task or at rest. FDG-fPET has most often been applied in simultaneous BOLD-fMRI/FDG-fPET (blood oxygenation level dependent functional MRI fluorodeoxyglucose functional positron emission tomography) imaging. BOLD-fMRI/FDG-fPET provides the capability to image the two primary sources of energetic dynamics in the brain, the cerebrovascular haemodynamic response and cerebral glucose uptake.

**Findings:** In this Data Note, we describe an open access dataset, Monash DaCRA fPET-fMRI, which contrasts three radiotracer administration protocols for FDG-fPET: bolus, constant infusion, and hybrid bolus/infusion. Participants (n=5 in each group) were randomly assigned to each radiotracer administration protocol and underwent simultaneous BOLD-fMRI/FDG-fPET scanning while viewing a flickering checkerboard. The Bolus group received the full FDG dose in a standard bolus administration; the Infusion group received the full FDG dose as a slow infusion over the duration of the scan, and the Bolus-Infusion group received 50% of the FDG dose as bolus and 50% as constant infusion. We validate the dataset by contrasting plasma radioactivity, grey matter mean uptake, and task-related activity in the visual cortex.

**Conclusions:** The Monash DaCRA fPET-fMRI dataset provides significant re-use value for researchers interested in the comparison of signal dynamics in fPET, and its relationship with fMRI task-evoked activity.

## Context

The neural functions of the human brain rely upon a stable and reliable energy supply delivered in the form of glucose[1]. The human brain accounts for 20% of the body’s energy consumption at rest[2],[3], of which 70-80% is used by neurons during synaptic transmission. Global and regional variations in the glucose uptake during neural activity can be measured using the [18]-fluorodeoxyglucose positron emission tomography (FDG-PET) method. As cerebral glucose uptake primarily reflects synaptic transmission[2], FDG-PET has long been used in neuroimaging studies as a proxy for neuronal activity. In recent years, functional brain imaging studies using the FDG-PET method have been somewhat overshadowed by the blood oxygenation level dependent functional magnetic resonance imaging (BOLD-fMRI) method. This is primarily due to the improved spatial and temporal resolution of fMRI in comparison to FDG-PET. Traditional FDG-PET methods provided a snapshot of glucose uptake averaged across the uptake and scan periods (approximate duration 30 mins), and were unable to distinguish between neural responses to stimuli presented closely in time. However, the recent availability of molecular MRI scanners which provide the capacity to simultaneously acquire BOLD-fMRI and FDG-PET data, has driven significant advances in FDG-PET methodologies for human neuroscience functional brain mapping studies [4–7].

Recently, improvements in radiotracer delivery have resulted in substantial improvement in the temporal resolution of FDG-PET. The method described as ‘functional’ PET (fPET) involves delivering the radiotracer as a constant infusion over the course of the scan. In a landmark study, Villien et al.[7] adapted the constant infusion technique[8] to deliver sufficient radiotracer to measure dynamic changes in brain glucose metabolism in response to a checkerboard stimulation, with a temporal resolution of 1-minute. Using fPET data acquired simultaneously with (non-functional) MRI (i.e., MRI/fPET), Villien et al. was able to estimate a general linear model response for blocked stimuli presented 5-10mins apart. Subsequent studies have extended these findings, and achieved fPET temporal resolutions of 1-minute[6,7,9–11] or less (12sec[5]; 16sec[12–14]; 30sec[15]).

The PET image quality relies upon the neural tissue radioactivity count rate from the administered radiotracer, and the duration of the scan[16]. fPET protocols typically have lower signal-to-noise ratio than static FDG-PET acquisitions, since the constant infusion approach must administer the same effective dose of radioactivity over a longer time period. Furthermore, fPET protocols require commencement of the scanning to be synchronised with the start of the radiotracer administration, to ensure that the measured brain activity is specific to the activity evoked during the experiment. Consequently, a constant infusion fPET scan has very little (close to zero) signal at the commencement of the experimental protocol, and the signal continuously increases over the duration of the infusion and scan[5,12]. Constant infusion fPET imaging protocols therefore tend to be quite long in comparison to standard FDG-PET and fMRI neuroimaging studies – usually around 90-100minutes[6,15]. These considerations restrict fPET studies primarily to populations that are able to be compliant with scanning requirements (e.g., restricted movement) over a long period of time.

The aim in acquiring the Monash <u>da</u>taset for <u>c</u>omparison of <u>r</u>adiotracer <u>a</u>dministration fPET-fMRI (‘**Monash DaCRA fPET-fMRI**’) was to contrast different radiotracer administration protocols for fPET data acquisition. The majority of fPET studies have used a constant infusion delivery protocol[6,7,9–11,14], where the entire dose of radiation is provided as an infusion over the course of the scan. However, a small number of studies have examined whether a hybrid bolus plus infusion protocol (bolus-infusion) might provide better signal at early timepoints while still allowing task-related activity to be measured at later timepoints. In a proof-of-concept comparison, we[12] found that a bolus-infusion protocol – where 50% of the dose was delivered as bolus, 50% as infusion – appeared to provide the most stable fPET signal for the longest period of time, compared to 100% constant infusion or 100% bolus protocols. Note however, this result was obtained in a case study design. Rischka et al.[5] used a 20% bolus plus 80% infusion protocol to test the lowest task duration detectable with fPET methodology. They were able to measure task-related activity (finger tapping) to stimuli separated by 2-min with an fPET frame size of 12-sec using this protocol; no signal was detected for stimuli separated by 1-min with 6-sec PET frames size. Rischka et al. concluded that the bolus-infusion protocol allowed assessment of reduced duration task blocks. However, they did not compare bolus-infusion to either constant infusion or bolus administration. In a subsequent study from the same group, Riscka et al.[17] demonstrated excellent reliability of fPET at rest with 20% bolus 80% infusion administration; reliability of fPET reduced during task performance; and was lowest for BOLD-fMRI during rest and task.

Here, we acquired fPET data with 50/50 bolus-infusion, 100% constant infusion and 100% bolus protocols. We chose to start with a proportional 50/50 bolus/infusion protocol, rather than some other fraction (e.g., 20/80, etc.) as a starting point for parsimony. Figure 1A illustrates our expectations for the fPET signal for the three protocols. Consistent with the results from our proof-of-concept case study, we expected that the bolus protocol would provide the largest overall signal magnitude, with the peak early in the scan period, decreasing in magnitude across the duration of the scan. The fPET signal for the constant infusion protocol was predicted to increase slowly through the course of the scan, with the overall lowest peak magnitude. Lastly, the fPET signal for the hybrid bolus-infusion protocol was expected to show the overall longest sustained period over the course of the scan. We predicted that the bolus-infusion protocol would provide the best sensitivity for detecting task-related effects in the checkerboard stimulus task, followed by the constant infusion then bolus protocol.

**Figure 1.**
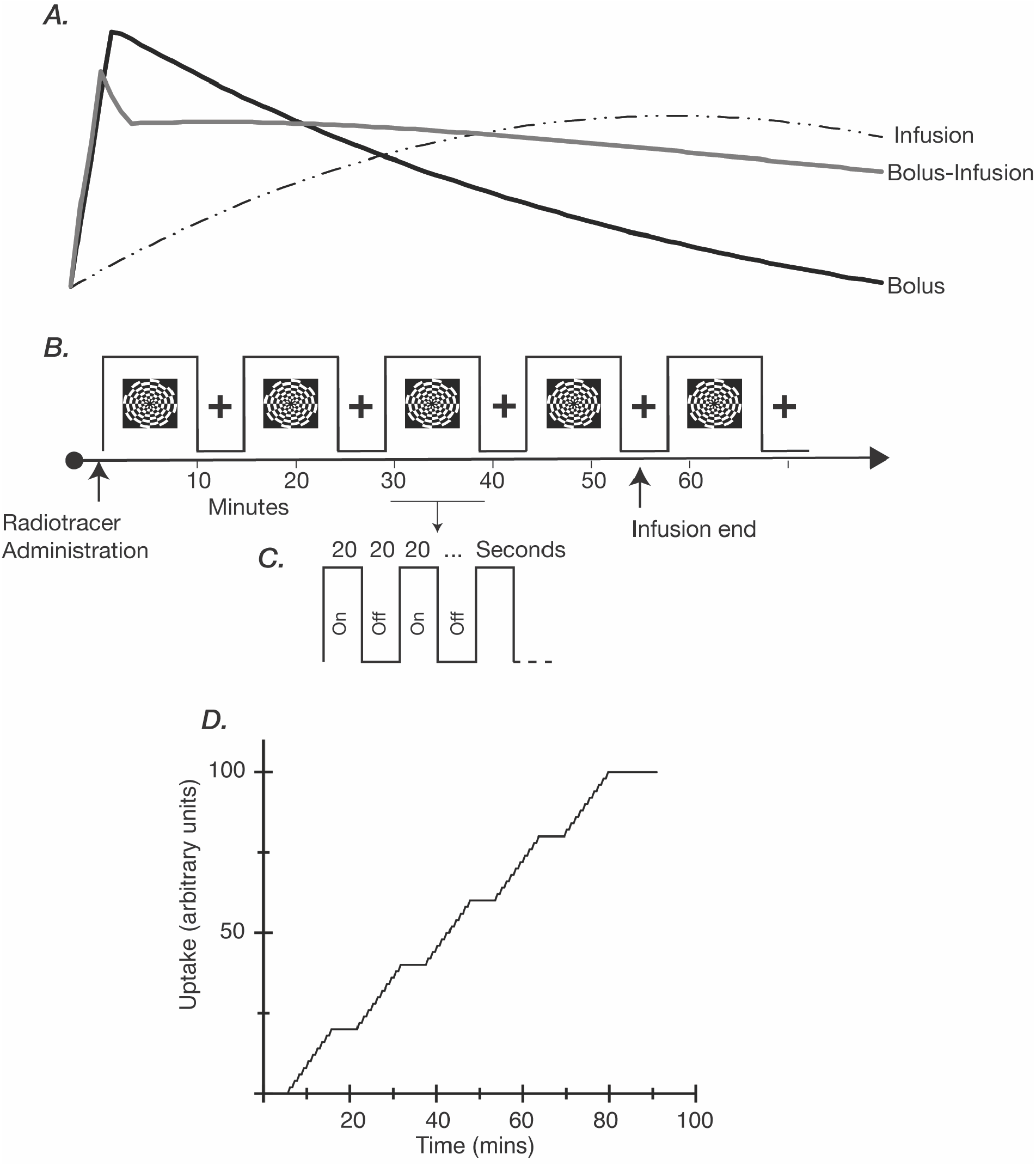
**A:** Hypothesised plasma radioactivity curves for the three administration protocols. Timing (i.e., signal peak and duration) is shown in comparison to the timing of the experimental protocol shown in panel B. We hypothesised that the bolus protocol would peak soon after administration, and decline rather quickly thereafter, returning to baseline levels by the end of the scan. We predicted that the bolus protocol would show the largest overall peak signal. For the infusion protocol we hypothesised that radioactivity would be close to zero at the beginning of the scan, continuing to rise for the duration of the scan. For the bolus-infusion protocol, we predicted that the peak signal would occur around the same time as the bolus protocol, but of smaller magnitude. We expected the signal would decrease slightly, but then remain at elevated levels for the duration of the scan. **B, C**. Experimental protocol. Checkerboard stimuli were presented in an embedded block design, with fast on/off periods (panel C) embedded within the longer ‘on’ (panel B) periods. **D**. Predicted task-related timecourse for the fPET general linear model.

We present one approach for GLM-based analysis of fPET data for data validation and quality control, and as an example of the type of analyses that are possible with this dataset. Development of more sophisticated methods of GLM and ICA analyses are examples of potential reuses of the dataset.

## Methods

All methods were reviewed by the Monash University Human Research Ethics Committee, in accordance with the Australian National Statement on Ethical Conduct in Human Research (2007). Administration of ionising radiation was approved by the Monash Health Principal Medical Physicist, in accordance with the Australian Radiation Protection and Nuclear Safety Agency Code of Practice (2005). For participants aged over 18-years, the annual radiation exposure limit of 5mSv applies; the effective dose in this study was 4.9mSv. Detailed information on the method for acquiring fPET data using bolus, constant infusion and bolus-infusion protocols is reported in Jamadar et al.[12].

### Available Data

Data is available on OpenNeuro with the accession number ds003397[18].

The data (Table 1) includes participant information (demography), scan information (e.g., start times), bloods (plasma radioactivity), raw MRI data (T1, T2 FLAIR, MR attenuation correction, susceptibility weighted images, field maps), unreconstructed PET data, and reconstructed PET images with temporal bins of 16sec.

**Table 1:**
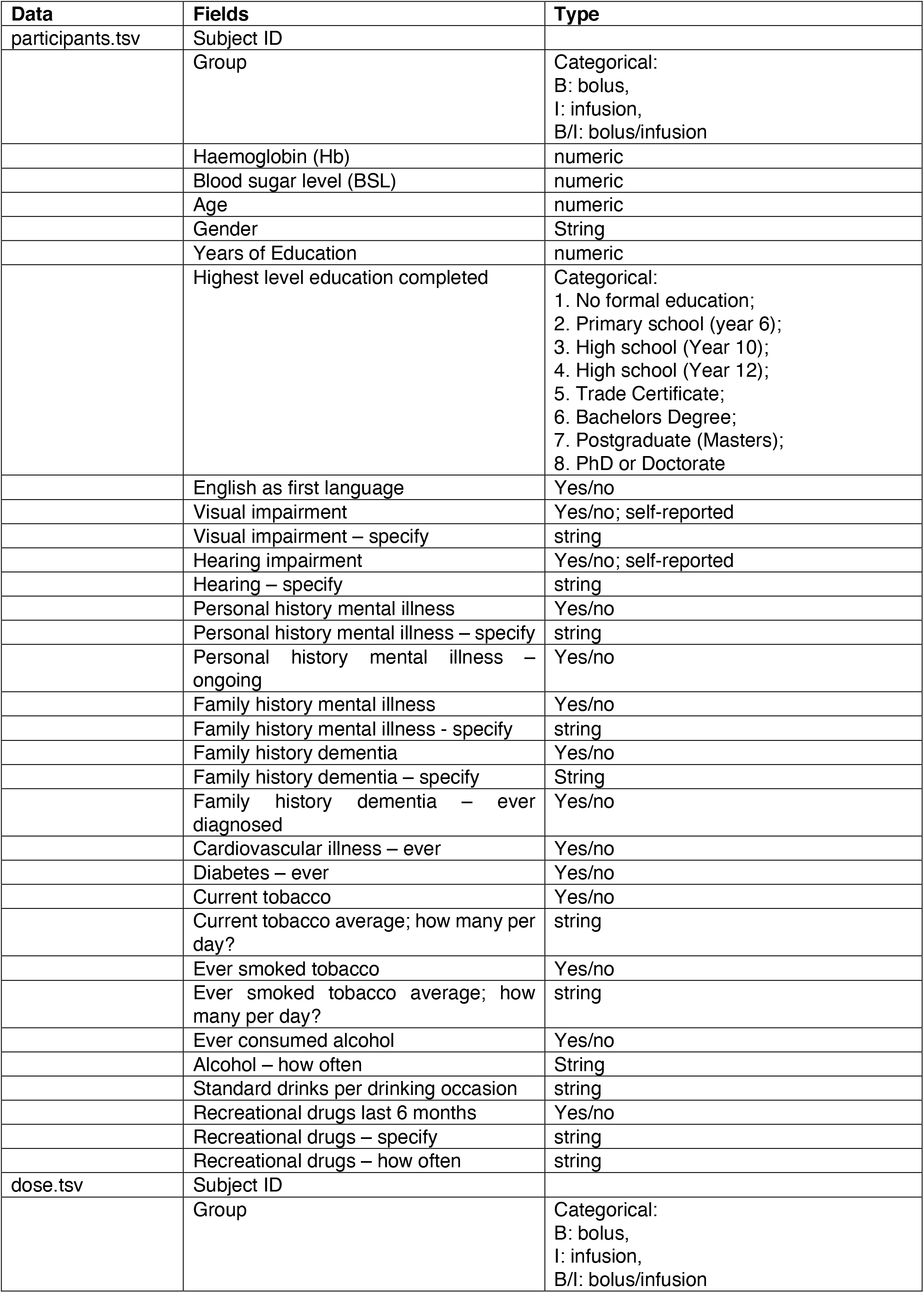
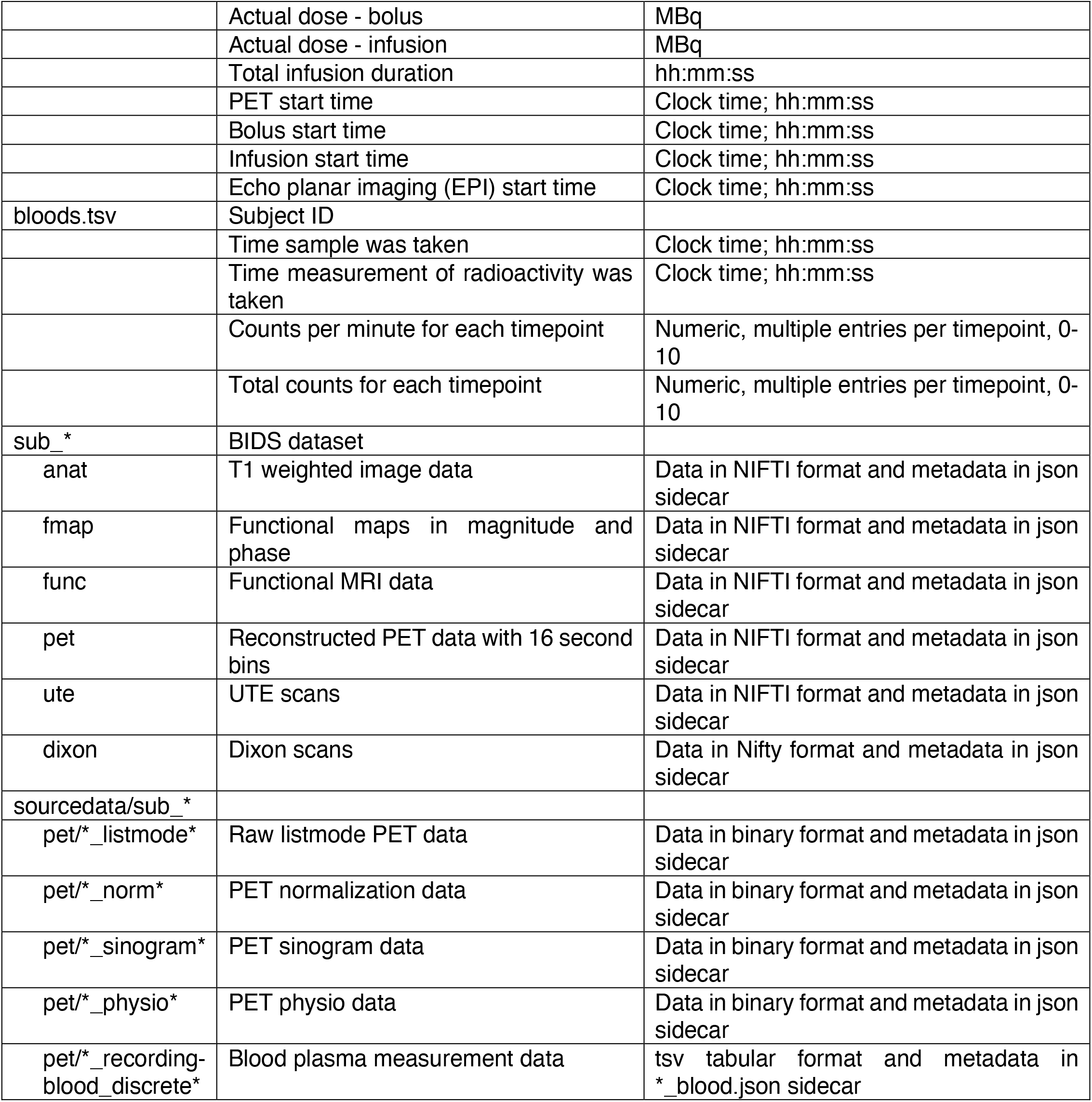
Data fields for the Monash radfPET-fMRI dataset

| Data | Fields | Type |
| --- | --- | --- |
| participants.tsv | Subject ID |  |
|  | Group | Categorical:<br>B: bolus,<br>I: infusion,<br>B/I: bolus/infusion |
|  | Haemoglobin (Hb) | numeric |
|  | Blood sugar level (BSL) | numeric |
|  | Age | numeric |
|  | Gender | String |
|  | Years of Education | numeric |
|  | Highest level education completed | Categorical:<br>1. No formal education;<br>2. Primary school (year 6);<br>3. High school (Year 10);<br>4. High school (Year 12);<br>5. Trade Certificate;<br>6. Bachelors Degree;<br>7. Postgraduate (Masters);<br>8. PhD or Doctorate |
|  | English as first language | Yes/no |
|  | Visual impairment | Yes/no; self-reported |
|  | Visual impairment – specify | string |
|  | Hearing impairment | Yes/no; self-reported |
|  | Hearing – specify | string |
|  | Personal history mental illness | Yes/no |
|  | Personal history mental illness – specify | string |
|  | Personal history mental illness – ongoing | Yes/no |
|  | Family history mental illness | Yes/no |
|  | Family history mental illness - specify | string |
|  | Family history dementia | Yes/no |
|  | Family history dementia – specify | String |
|  | Family history dementia – ever diagnosed | Yes/no |
|  | Cardiovascular illness – ever | Yes/no |
|  | Diabetes – ever | Yes/no |
|  | Current tobacco | Yes/no |
|  | Current tobacco average; how many per day? | string |
|  | Ever smoked tobacco | Yes/no |
|  | Ever smoked tobacco average; how many per day? | string |
|  | Ever consumed alcohol | Yes/no |
|  | Alcohol – how often | String |
|  | Standard drinks per drinking occasion | string |
|  | Recreational drugs last 6 months | Yes/no |
|  | Recreational drugs – specify | string |
|  | Recreational drugs – how often | string |
| dose.tsv | Subject ID |  |
|  | Group | Categorical:<br>B: bolus,<br>I: infusion,<br>B/I: bolus/infusion |
|  | Actual dose - bolus | MBq |
|  | Actual dose - infusion | MBq |
|  | Total infusion duration | hh:mm:ss |
|  | PET start time | Clock time; hh:mm:ss |
|  | Bolus start time | Clock time; hh:mm:ss |
|  | Infusion start time | Clock time; hh:mm:ss |
|  | Echo planar imaging (EPI) start time | Clock time; hh:mm:ss |
| bloods.tsv | Subject ID |  |
|  | Time sample was taken | Clock time; hh:mm:ss |
|  | Time measurement of radioactivity was taken | Clock time; hh:mm:ss |
|  | Counts per minute for each timepoint | Numeric, multiple entries per timepoint, 0-10 |
|  | Total counts for each timepoint | Numeric, multiple entries per timepoint, 0-10 |
| sub_* | BIDS dataset |  |
| anat | T1 weighted image data | Data in NIFTI format and metadata in json sidecar |
| fmap | Functional maps in magnitude and phase | Data in NIFTI format and metadata in json sidecar |
| func | Functional MRI data | Data in NIFTI format and metadata in json sidecar |
| pet | Reconstructed PET data with 16 second bins | Data in NIFTI format and metadata in json sidecar |
| ute | UTE scans | Data in NIFTI format and metadata in json sidecar |
| dixon | Dixon scans | Data in Nifty format and metadata in json sidecar |
| sourcedata/sub_* |  |  |
| pet/*_listmode* | Raw listmode PET data | Data in binary format and metadata in json sidecar |
| pet/*_norm* | PET normalization data | Data in binary format and metadata in json sidecar |
| pet/*_sinogram* | PET sinogram data | Data in binary format and metadata in json sidecar |
| pet/*_physio* | PET physio data | Data in binary format and metadata in json sidecar |
| pet/*_recording-blood_discrete* | Blood plasma measurement data | tsv tabular format and metadata in *_blood.json sidecar |

### Participants

Fifteen young adults participated in this study. Participants were randomly assigned to the bolus, infusion, and bolus-infusion groups. Participants were aged 18-20 years, right handed, normal or corrected-to-normal vision, and were screened for diabetes, hearing impairment, personal or family history of mental or neurodegenerative illness, and personal history of head injury or neurological condition. Women were screened for pregnancy. Prior to the scan, participants were directed to consume a high protein/low sugar diet for 24-hrs, fast for 6-hrs, and drink 2-6 glasses of water.

Participants in the bolus group had mean age 19.2 years, three were male, and had 12-14 years of education (mean 13.2 years). Participants in the infusion group had mean age 19.4 years, four were male, and had 14-15 years of education (mean 14.6 years). Participants in the bolus-infusion group had mean age 19.4 years, one was male, and had 12-15 years of education (mean 13.6 years).

### Stimuli and Tasks

Participants rested with eyes closed during the initial 20-mins while non-functional MR scans were acquired. During simultaneous fMRI-fPET scanning, participants viewed flickering checkerboard stimuli presented in an embedded block design[6]. We have previously shown that an embedded design provides simultaneous contrast for task-evoked BOLD-fMRI and FDG-fPET data. The task alternates between 640-sec flashing checkerboard blocks and 320-sec rest blocks (Figure 1B). This slow alternation provides fPET contrast. Within the 640-sec checkerboard blocks, checkerboard and rest period alternate with a rate of 20-sec on, 20-sec off (Figure 1C). This fast alternation is suitable for BOLD-fMRI contrast.

The checkerboard stimulus was a circular checkerboard of size 39cm (visual angle 9°) presented on a black background. The checkerboard flickered (i.e., alternated black and white segments) at 8Hz. During the ‘off’ periods, participants rested with eyes fixated on a white cross of size 3cm (visual angle (0° 45’).

### Procedure

Participants were cannulated in the vein in each forearm with a minimum size 22-gauge cannula. A 10mL baseline blood sample was taken at time of cannulation. For all participants, the left cannula was used for FDG administration, and the right cannula was used for blood sampling. Primed extension tubing was connected to the right cannula for blood sampling via a three way tap.

Participants underwent a 95-min simultaneous MRI-PET scan in a Siemens Biograph 3Tesla molecular MR (mMR) scanner. Participant lay supine in the scanner bore with head in a 16-channel radiofrequency head coil, and were instructed to lie as still as possible. [18F]-FDG (average dose = 238MBq), was administered either as a bolus, an infusion, or as a bolus-infusion (50% bolus 50% infusion). For the infusion protocols, infusion rate was 36mL/hr using a BodyGuard 323 MR-compatible infusion pump (Caesarea Medical Electronics, Caesarea, Israel). For the bolus protocol, the bolus was administered at the time of the PET scan onset. For the infusion protocol, the infusion commenced at the time of PET scan onset. For the bolus-infusion protocol, the bolus was administered at the onset time of the PET scan, and the infusion started as soon as possible (average = 40-sec) after the bolus. For the infusion and bolus-infusion protocols, the infusion ceased at 55-mins. We hypothesised that the plasma radioactivity would be maintained for a short period thereafter, however this was not the case (see Results Section 3.1).

Plasma radioactivity levels were measured throughout the duration of the scan. At 5-mins post-administration, a 10mL blood sample was taken from the right forearm using a vacutainer; the time of the 5mL mark was noted for subsequent decay correction. Subsequent blood samples were taken at 5-min intervals. The cannula line was flushed with 10mL of saline after every sample to minimise line clotting. Immediately after sampling, the sample was placed in a Hereaus Megafuge 16 centrifuge (ThermoFisher Scientific, Osterode, Germany) and spun at 2000rpm for 5-mins; 1000μL was pipetted, transferred to a counting tube, and placed in a well counter for 4-mins. The count start time, total number of counts, and counts per minute were recorded for each sample.

### MR-PET Protocol

PET data (90:56-min) was acquired in list mode. The onset of the PET acquisition (and the radiotracer administration) was locked to the onset of the T2* EPIs.

The MRI and PET scans were acquired in the following order: (i) T1-weighted 3D MPRAGE (TA = 7.01 min, TR = 1,640 ms, TE = 2.34 ms, flip angle = 8°, FOV = 256 × 256 mm2, voxel size = 1 × 1 x 1 mm3, 176 slices, sagittal acquisition; (ii) T2-weighted FLAIR (TA = 5.78 min); (iii) SWI (TA = 6.83 min); (iv) gradient field map TA = 1.08 min; (v) MR attenuation correction Dixon (TA = 0.65 min, TR = 4.14 ms, TEin phase = 2.51 ms, TE out phase = 1.3 ms, flip angle = 10°); (vi) T2*-weighted echo-planar images (EPIs) (TA = 90:56 min; TR=4000ms, TE=30ms, FOV=190mm, 3×3×3mm voxels, 44 slices, ascending axial acquisition), P-A phase correction (TA = 0.37 min); (vii) UTE (TA = 1.97 min).

## Data Records

Detailed information about the data records available for the Monash DaCRA fPET-fMRI dataset (OpenNeuro ds003397)[18] is reported in Table 1. Table 2 reports the software used in this manuscript.

**Table 2:**
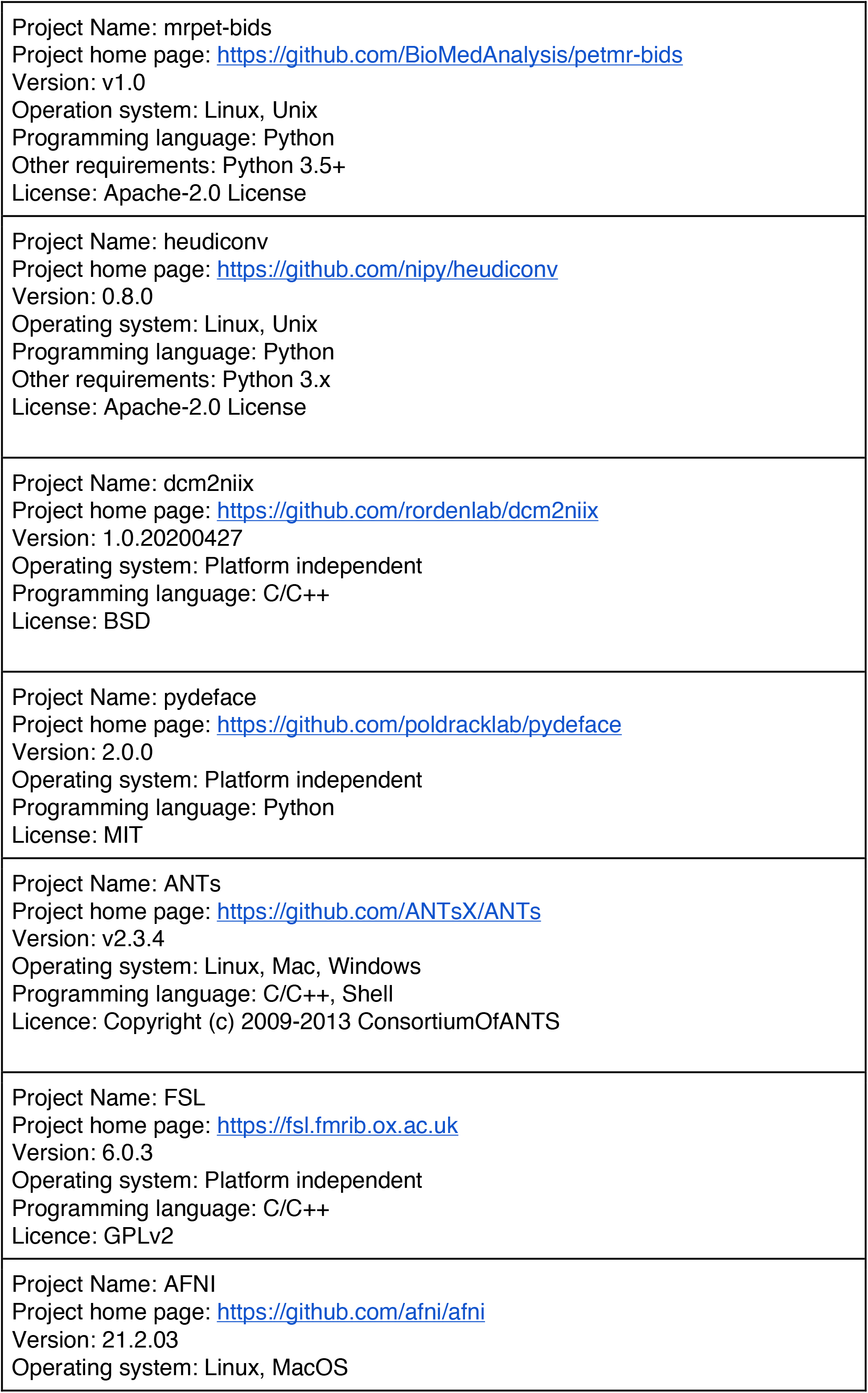
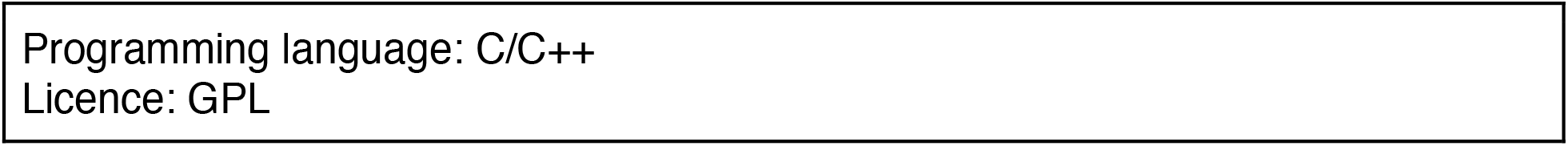
Software used in the development of this manuscript.

|  |
| --- |
| Project Name: mrpet-bids<br>Project home page: https://github.com/BioMedAnalysis/petmr-bids<br>Version: v1.0<br>Operation system: Linux, Unix<br>Programming language: Python<br>Other requirements: Python 3.5+<br>License: Apache-2.0 License |
| Project Name: heudiconv<br>Project home page: https://github.com/nipy/heudiconv<br>Version: 0.8.0<br>Operating system: Linux, Unix<br>Programming language: Python<br>Other requirements: Python 3.x<br>License: Apache-2.0 License |
| Project Name: dcm2niix<br>Project home page: https://github.com/rordenlab/dcm2niix<br>Version: 1.0.20200427<br>Operating system: Platform independent<br>Programming language: C/C++<br>License: BSD |
| Project Name: pydeface<br>Project home page: https://github.com/poldracklab/pydeface<br>Version: 2.0.0<br>Operating system: Platform independent<br>Programming language: Python<br>License: MIT |
| Project Name: ANTs<br>Project home page: https://github.com/ANTsX/ANTs<br>Version: v2.3.4<br>Operating system: Linux, Mac, Windows<br>Programming language: C/C++, Shell<br>Licence: Copyright (c) 2009-2013 ConsortiumOfANTS |
| Project Name: FSL<br>Project home page: https://fsl.fmrib.ox.ac.uk<br>Version: 6.0.3<br>Operating system: Platform independent<br>Programming language: C/C++<br>Licence: GPLv2 |
| Project Name: AFNI<br>Project home page: https://github.com/afni/afni<br>Version: 21.2.03<br>Operating system: Linux, MacOS |

Participants.tsv is a text file reporting demographic and anthropometric data for each subject, ordered by subject ID. Plasma_radioactivity.tsv is a text file reporting the plasma radioactivity counts and measurement times for each subject, ordered by subject ID.

The dataset contains both raw (unprocessed) images and source data (i.e., unreconstructed PET listmode data). Both are organised in sub-directories that correspond to subject ID, according to BIDS (for MRI) or BIDS-consistent (for PET) specification. For each subject, T1-weighted MPRAGE images, fMRI images, and gradient field maps are in the *anat* (anatomical data), *func* (functional MRI data) and *fmap* (field map) subdirectories, along with metadata in the json sidecar. Dixon and UTE scans are available for PET source data reconstruction, which are organised into *dixon* and *ute* sub-directories.

Although there is not currently a listmode PET BIDS specification, the same structure is followed with a json sidecar accompanying the image data. PET image data was obtained by reconstructing the PET source data into 16-sec bins offline using Siemens Syngo E11p. Attenuation was corrected using pseudoCT[19] Ordinary Poisson-Ordered Subset Expectation Maximisation (OP-OSEM) algorithm with point spread function modelling[20] with 3 iterations, 21 subsets and 344×344×127 (voxel size = 2.09×2.09×2.03mm^3^) reconstruction matrix size. A 5mm 3D Gaussian post-filtering was applied to the final reconstructed images. Following the BIDS extension for PET (BEP009), blood data are also included in the *pet* directory, which report the plasma radioactivity counts and measurement times for the subject. Data in sub-*/dixon, sub-*/ute and sub-*/pet are ignored in the BIDS validation process, as they are not officially supported by the current BIDS specification.

The *sourcedata* directory contains the raw, un-reconstructed PET source data that was directly exported from the Siemens scanner console. The source data includes PET listmode data, normalisation data, sinogram data, and physiology data. The raw PET data are in the form of a file pair (one DICOM header and one binary file) with the two paired files having the same file name but different extensions (.dcm for DICOM, .bf for binary). A json metadata sidecar file was added to each subject’s raw dataset, consistent with the BIDS approach for supported structures. The blood plasma radioactivity data is included and is identical to the reconstructed PET image data. The *sourcedata* directory is also excluded in the BIDS validation process.

To prepare the BIDS dataset, the open source conversion tool *Heudiconv* (https://github.com/nipy/heudiconv) was used to organise the imaging data into structured directory layouts, and the dcm2niix converter (https://github.com/rordenlab/dcm2niix) was used to convert image data from dicom to nifti format. Following the approach in our previous manuscript[21], we applied scripts to: (i) remove personal identifiable information from the raw PET dicom header; (ii) add custom json sidecar files to the PET raw data and reconstructed image data; and (iii) generate plasma radioactivity files. Refer to https://github.com/BioMedAnalysis/petmr-bids for these scripts.

Defacing was applied to T1-weighted, Dixon and UTE images using pydeface (https://github.com/poldracklab/pydeface). Reconstructed PET images and PET raw data were not defaced as subjects cannot be visually identified from the PET images.

## Data Validation - Methods

We validated the Monash DaCRA fPET-fMRI dataset by confirming that the data yielded expected results with standard general linear model analysis.

### fMRI Image Preparation and Analysis

The subjects’ T1 brain images were extracted (ANTs[22]), to standard space using affine transformation (12 degrees of freedom) and a standard space 2mm brain atlas. The EPI scans for all subjects underwent a standard fMRI pre-processing pipeline. All EPI scans were brain extracted (FSL BET[23]), corrected for intensity nonuniformity using N4 Bias field correction (ANTs[24]), motion corrected (FSL MCFLIRT[25]), and slice timing corrected (AFNI).

Pre-processed fMRI data was submitted to a subject-level GLM using FSL[26] FEAT. The following pre-statistics were applied: spatial smoothing using a Gaussian kernel of FWHM 5mm; grand-mean intensity normalisation of the entire 4D dataset by a single multiplicative factor; and highpass temporal filtering (Gaussian-weighted least-squares straight line fitting, with sigma=50.0s). Time-series statistical analysis was carried out using FILM with local autocorrelation correction[27]. For the subject-level analysis we used a GLM where the only regressor of interest was task, and temporal derivative as covariate. Subject-level Z (Gaussianised) static images were thresholded non-parametrically using clusters determined by Z > 1.6 and a corrected cluster significance threshold of P=0.05[28]. Group-level analysis was carried out using FLAME (FMRIB’s Local Analysis of Mixed Effects) stage 1[29–31] to obtain the group mean. Three separate group-level GLMs were conducted for each group (bolus, infusion, bolus-infusion).

### PET Image Preparation and Analysis

Spatial realignment was performed on the dynamic FDG-fPET images using FSL MCFLIRT[25]. A mean FDG-PET image was derived from the entire dynamic timeseries and rigidly normalized to the individual’s high-resolution T1-weighted image using ANTs[22]. The dynamic FDG-fPET images were then normalized to MNI space using the rigid transform in combination with the non-linear T1 to MNI warp. fPET images were spatially smoothed using a Gaussian kernel of 12mm FWHM.

fPET data processing was carried out using FEAT (fMRI Expert Analysis Tool) version 6.00. The pre-processed smoothed MNI152-space fPET images were submitted to a GLM analysis using FILM[27].

We modelled the increasing whole brain radioactivity signal related to radiotracer uptake across the PET scan period. For each subject, we assume an underlying baseline activity (Y_base_) when no task was performed to model the radiotracer uptake. We subtracted Y_base_ from the timeseries data to obtain baseline corrected data:

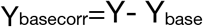

where Y_base_ is the underlying baseline timeseries for each voxel. Then, we can estimate β_task_ as

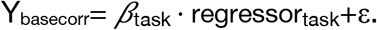

In another GLM, we approximate the baseline using grey matter mean (regressor_GM_) as confound:

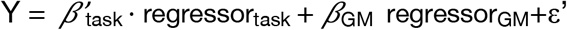

Thus, the ‘cleaned’ data is represented as

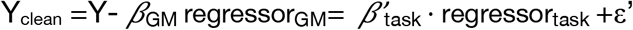

also

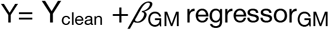

Since

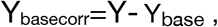

Then,

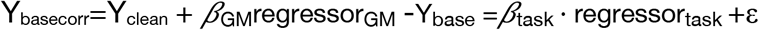

i.e.

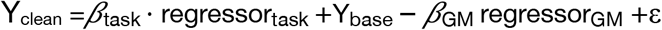

Since the baseline Y_base_ coefficient of 1 can be expressed as

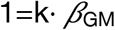

then

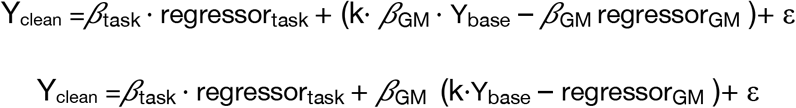

Hahn et al.[10] reported that the change in the baseline signal in grey matter ROIs during the scan period was approximately linear. We assumed that the grey matter mean signal over the scanning period showed a similar trend (see our data in Fig 2B) and therefore used a linear change n·regressor_line_ to approximately replace (k·Y_base_ – regressor_GM_).

**Figure 2:**
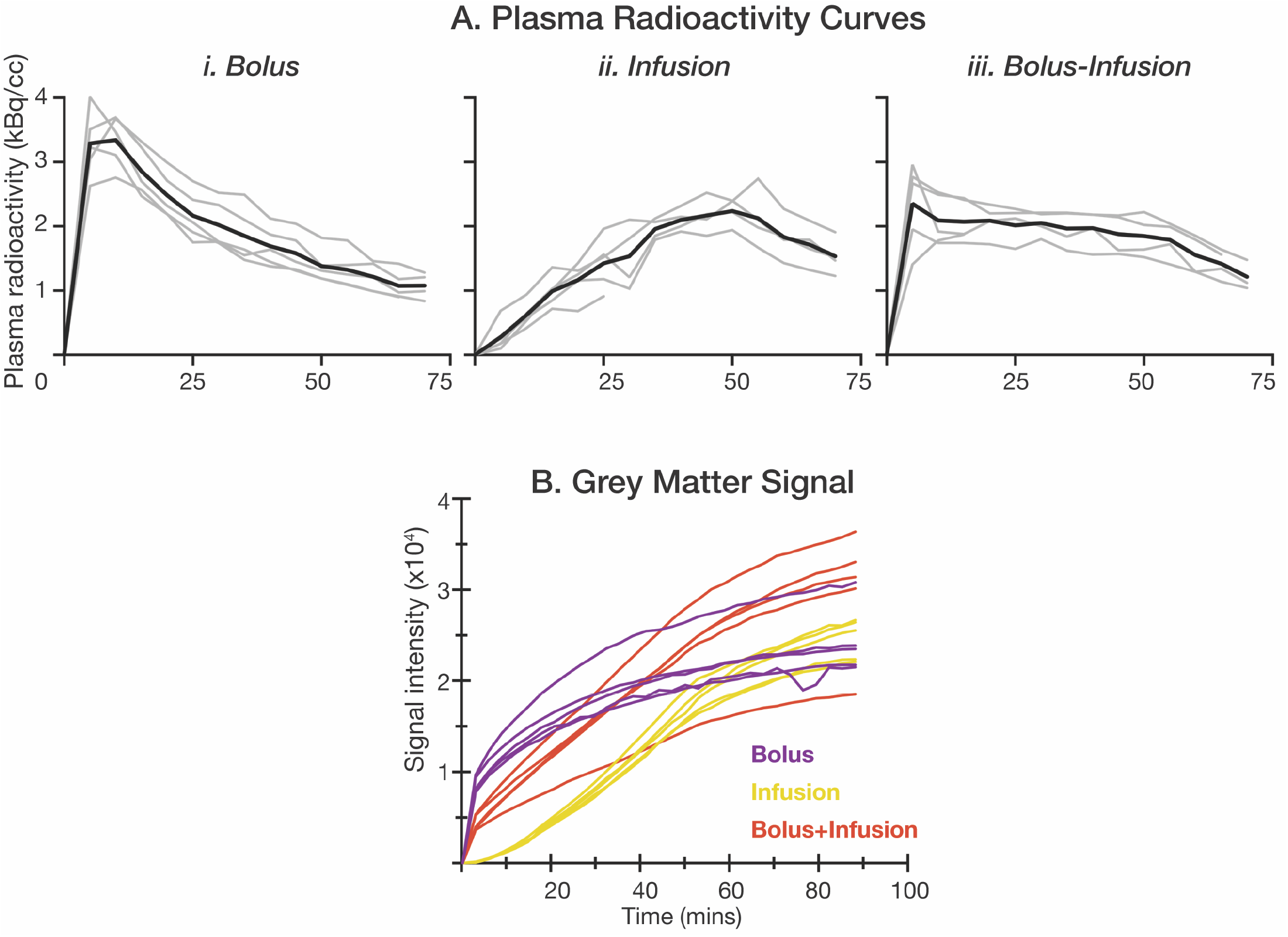
**A**. Plasma radioactivity curves for **i**. bolus administration, **ii**. infusion administration and **iii**. bolus-infusion protocol. Black line shows average radioactivity and grey lines show activity for individual subjects. **B**. Average grey matter signal across all voxels for each subject.

Then

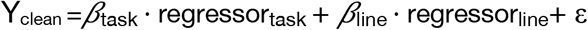

where

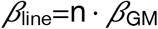

The subject-level GLM had two regressors: namely a task regressor (Figure 1D) and a linear regressor that modelled the continuous underlying baseline uptake over time.

Subject-level Z (Gaussianised) static images were thresholded non-parametrically using clusters determined by Z > 1.6 and a corrected cluster significance threshold of P=0.05[28]. Group-level analysis was carried out using FLAME (FMRIB’s Local Analysis of Mixed Effects) stage 1[29–31] to obtain the group mean activation map. Since the baseline uptake rate differed throughout the brain, in some voxels the baseline regressed timeseries data showed a negative trend because the uptake rates was lower than the grey matter mean uptake rate. To determine the brain regions that associated negatively with the task we included a regressor to model negative task events.

## Data Validation -Results

### Plasma Radioactivity

We hypothesised shapes of the radioactivity curves for the three groups assuming that the bolus-infusion protocol would provide the best sensitivity for detecting task-related effects in the checkerboard stimulus task, followed by the constant infusion and the bolus protocol (Figure 1A). The measured plasma radioactivity curves for the three radiotracer administration protocols are shown in Figure 2A. Radioactivity peaked early and declined quickly for the bolus protocol. The largest radioactivity peak was evident in the bolus protocol. In the infusion protocol, radioactivity continued to rise until the cessation of the infusion (55-mins), at which point activity declined. The continued upward slope of the curved for the duration of the infusion suggests that the plasma radioactivity had not yet reached its peak before the cessation of the infusion. As predicted, the bolus-infusion protocol showed an early peak after the bolus; the activity decreased slightly but was maintained at close to a constant level for the duration of the infusion. As expected, the peak for the bolus-infusion protocol was smaller than in the bolus protocol.

As noted in the methods, for the infusion and bolus-infusion protocols we ceased infusion at the 55-min mark. We expected that radioactivity would remain stable for a short period of time afterwards. However, both protocols showed a clear decline in radioactivity when infusion ceased.

In sum, on the basis of the plasma radioactivity curves alone, it is apparent that the bolus-infusion protocol provides the most stable signal over the course of the scan, which is maintained as long as infusion is administered.

### Grey Matter Signal

Consistent with the plasma radioactivity results, the grey matter mean signal increased fastest for bolus administration, followed by bolus-infusion, with the infusion protocol showing the slowest increase in signal (Fig. 2B). By the end of the experiment, four out of the five bolus-infusion subjects showed the highest signal intensity, with most (4/5) bolus subjects showing a similar level of signal intensity to the infusion only subjects.

### fMRI Results

The fMRI results are shown primarily to confirm that the experimental design was successful in eliciting stimulus-evoked fMRI responses in the visual cortex (Fig. 3). As expected, visual cortex was active for all three groups (and in the average across the fifteen subjects; Fig. 3D); additional activity was also apparent in other cortical areas known to be involved in processing visual stimuli, including the intraparietal sulcus and frontal eye fields.

**Figure 3:**
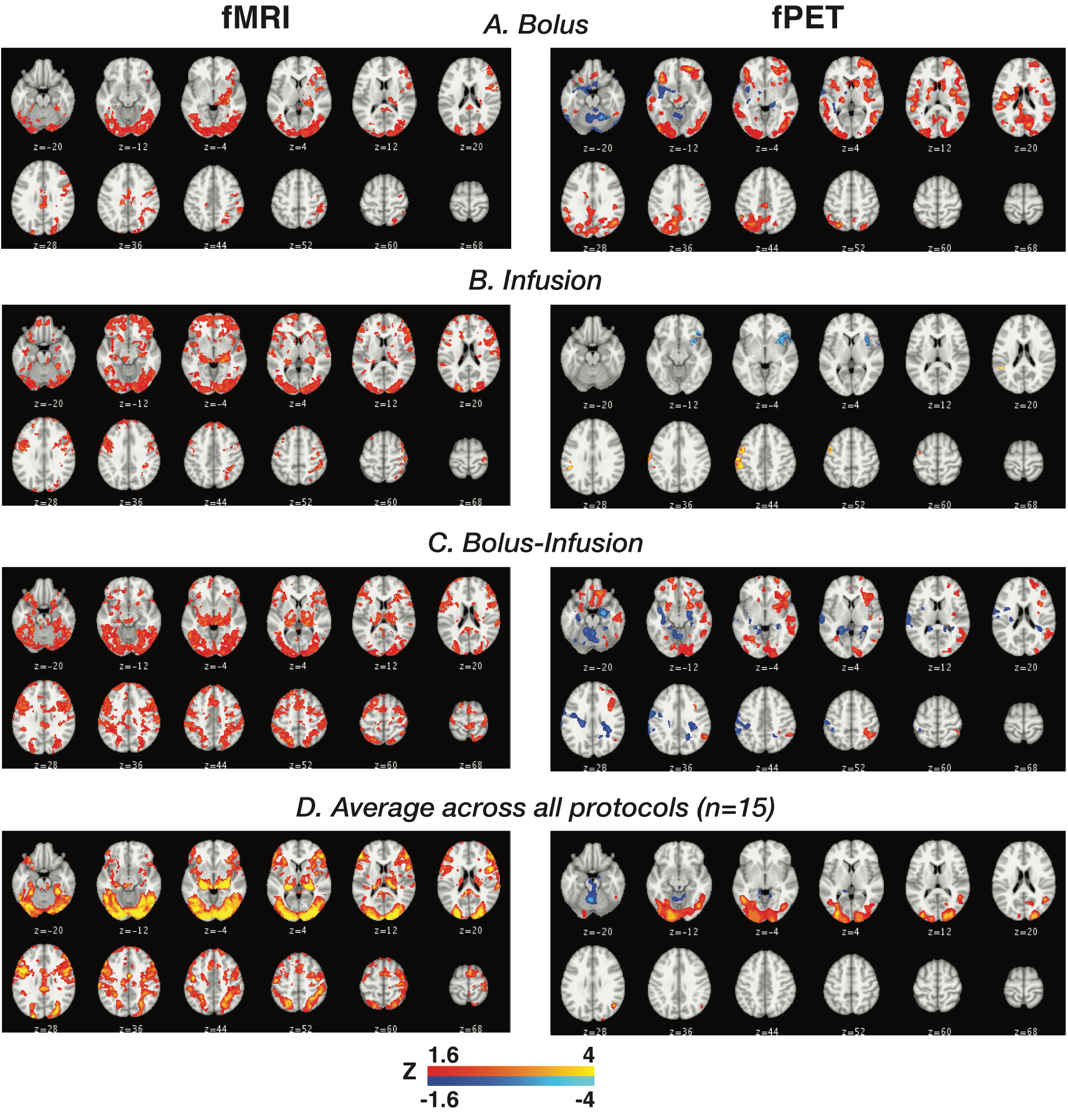
Group-level activation maps for task (Zcorr>1.6) for (left) fMRI and (right) fPET; shown separately for **A** bolus group, **B** infusion group, **C** bolus-infusion group. Given that the fMRI protocol did not differ for the three groups we also show the group average fMRI across all fifteen subjects in panel **D**.

### fPET Results

Across the three protocols (Fig. 3) task-related fPET showed a more focal pattern of activity in the visual cortex compared to fMRI. Visual comparison of the three administration protocols showed only modest levels of activity in the infusion-only protocol (Fig. 3B), with more widespread cortical activity in the bolus-only (Fig. 3A) and bolus-infusion protocols (Fig. 3C). The bolus-infusion protocol showed more widespread ‘negative’ uptake than the other administration protocols, suggesting that these regions showed slower uptake of FDG by comparison to the grey matter mean.

We visualised individual variability in percent signal change in five regions of interest for fMRI and fPET (Fig. 4). ROIs were defined as those that showed suprathreshold activity -2.3> *z* >2.3 in the middle blocks (blocks 2, 3, 4) of the fPET data. Blocks 2,3,4 were chosen to coincide with the most stable activity across the three administration protocols. Figure 4 (right panels) shows the regions of interest. In the primary visual cortex (Fig. 4A, B), subjects uniformly showed positive percent signal change for both the fMRI and fPET. In the frontal regions of interest, fPET showed a uniform negative percent signal change, suggesting slower uptake compared to grey matter; whereas fMRI showed close to zero percent signal change for all subjects. It is notable that within each group (bolus, infusion, bolus-infusion) there is quite a bit of variability between individuals of 1-1.5% for both fMRI and fPET. Evaluating the fPET percent signal change, no single administration method appears to provide a more consistent fPET signal change across individuals; or a higher fPET signal change than the others.

**Figure 4.**
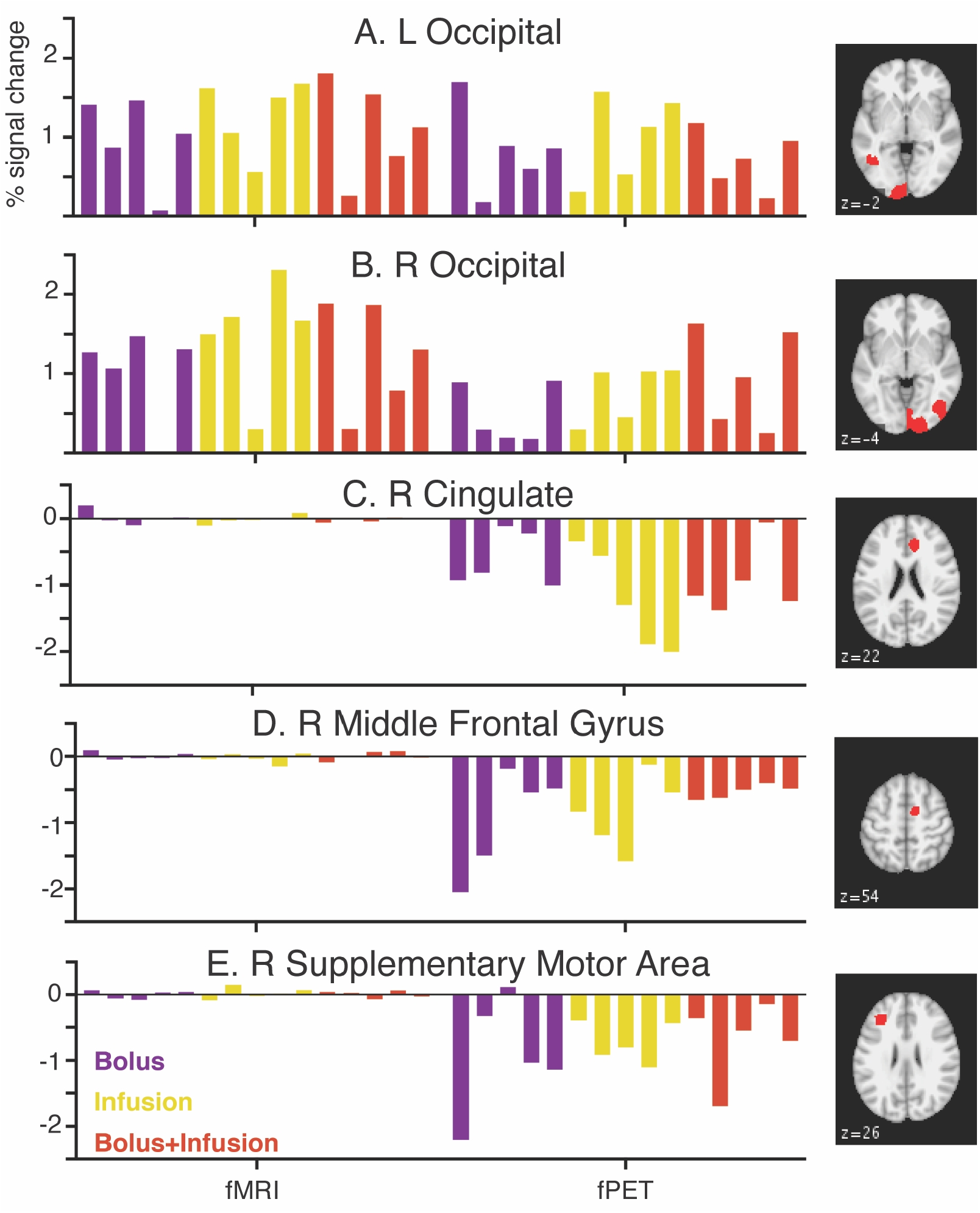
Percent signal change for five regions of interest for each administration method. Each column represents a single subject. Abbreviations: L, left; R, right.

Finally, since Figure 2 suggests that each administration protocol shows different timeframes for peak signal, we visualised fPET activity across three blocks at the start (blocks 1,2,3), middle (blocks 2,3,4) and end (blocks 3,4,5) of the scan period (Fig. 5). The bolus only protocol (Fig. 5A) showed the largest amount of suprathreshold activity at the start of the experiment (blocks 1,2,3), with less activity in the middle and end of the experiment. While activity in the visual cortex is evident, there is substantial additional suprathreshold activity across the cortex. The infusion only protocol (Fig. 5B) showed the least amount of suprathreshold activity across the blocks. Even though signal uptake is highest at the end of the experiment for this protocol (Fig. 2), there is little suprathreshold activity in the visual cortex evident during this period (blocks 3,4,5). Suprathreshold visual cortex activity is evident in the middle blocks (2,3,4) for this protocol. The bolus-infusion protocol (Fig. 5C) showed the most sustained suprathreshold visual cortex activity compared to the bolus and infusion protocols; activity was evident in blocks 1,2,3 and 2,3,4; but little activity in blocks 3,4,5. Like the bolus only group, the bolus-infusion group showed additional activity outside the visual cortex, which may represent false positive activity.

**Figure 5.**
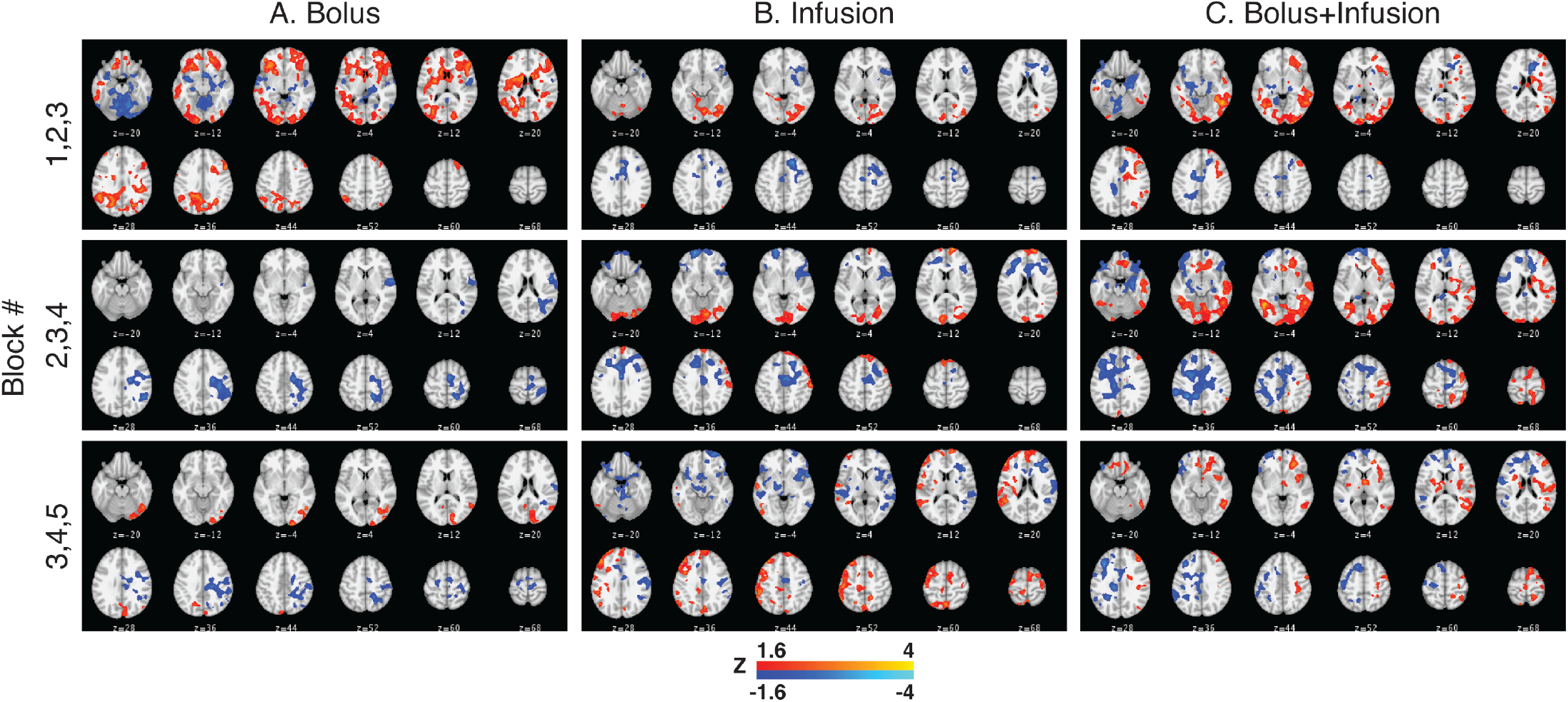
fPET results for blocks 1,2,3 (top), 2,3,4 (middle), 3,4,5 (bottom) for each administration protocol.

## Concluding Remarks and Re-use Potential

Simultaneous MR-PET is a nascent technique, opening up many opportunities for scientific discovery, methods development and signal optimisation of dual-modality data. Although very few imaging facilities world-wide currently possess the infrastructure and technical skill to acquire fPET-fMRI data, the rapid increase in publication (e.g.[5–7,10–12,14,15,21,32,33]) and reuse metrics of publicly available datasets[34,35] attests to the value the international neuroscience community places on this novel data type. The **Monash DaCRA fPET-fMRI** dataset is the only publicly available dataset that allows comparison of radiotracer administration protocols for fPET-fMRI. We provide both raw (listmode) and reconstructed fPET data to maximise the re-use value of the dataset. With listmode and reconstructed data, examples of re-use include the development of new processing pipelines and signal optimisation methods that take into account variability in radiotracer dynamics related to differences in administration method. Release of listmode PET data is notable, to our knowledge only one other open source dataset includes listmode PET data; the Monash visfPET-fMRI dataset[21]. These data releases are very novel, occurring prior to the formalisation of the PET BIDS standard (BEP009)[36,37]. The draft PET BIDS standard does not yet extend to listmode data[37], so we applied our BIDS-like standard[21] to ensure that it is consistent with the *Interoperable* principle of the FAIR philosophy. We[21] have previously demonstrated that listmode fPET data can be accurately reconstructed using open source methods STIR[38] and SIRF[39], confirming the Monash DaCRA fPET-fMRI dataset is also consistent with the *Reusable* principle of the FAIR philosophy.

Open source fPET-fMRI datasets provide many opportunities for progress in methods development: in the acquisition of the images, image reconstruction, data preparation and analysis. The complementary nature of haemodynamic and ‘glucodynamic’ responses to brain activity also presents an excellent opportunity for neuroscientific discovery. One area where further development is required is in the development of accurate general linear models (GLMs) for the analysis of task-based responses. Standard practices exist for GLM analysis of fMRI data (e.g., SPM & FSL-based approaches), but these do not yet exist for fPET data. The Vienna group[5,10,11,15] have reported a number of GLM-based analyses, which are analogous to block-design fMRI analyses. A number of questions remain: for example, there has not yet is not yet agreement in the best way to manage the increasing baseline signal related to radiotracer dynamics over the course of the scan. Here we have presented one approach to GLM analysis of task-based fPET data, however more work is required to validate the approach. This dataset provides an excellent opportunity develop task-based fPET analyses that account for underlying variability in radiotracer administration and uptake.

## List of Abbreviations

BIDS: Brain imaging data structure
BOLD-fMRI/FDG-fPET: Blood oxygenation level dependent functional magnetic resonance imaging [18F]-fluorodeoxyglucose functional positron emission tomography
DaCRA: Dataset for comparison of radiotracer administraion
EPI: Echo planar images
FDG: [18F]-fluorodeoxyglucose
FDG-*f*PET: [18F]-fluorodeoxyglucose functional positron emission tomographer
FDG-PET: [18F]-fluorodeoxyglucose positron emission tomography
FLAIR: Fluid attenuation inversion recovery
fMRI: Functional magnetic resonance imaging
FOV: Field of view
fPET: Functional positron emission tomography
fPET-fMRI: Simultaneous functional positron emission tomography functional magnetic resonance imaging
GLM: General linear model
ICA: Independent component analysis
ID: Identifier
MPRAGE: Magnetisation prepared rapid gradient echo
MRI: Magnetic resonance imaging
OP-OSEM: Ordinary Poisson ordered subset expectation maximisation
P-A: Posterior-anterior
PseudoCT: Pseudo computed tomography
ROI: Region of interest
SWI: Susceptibility weighted imaging
TA: Acquisition time
TE: Echo time
TR: Repetition time
UTE: Ultrashort echo time

## Consent for Publication

Not applicable

## Author Contributions

The research question was designed by SDJ and PGDW; the research design was developed by PGDW, AC, RM & SDJ. Funding was obtained by GFE, SDJ & ZC. Data was prepared for release by EXL, SZ, AC & RM. Manuscript was written by SDJ, EXL & SZ. All authors have contributed to manuscript preparation.

